# AAV-BR1 targets endothelial cells in the retina to reveal their morphological diversity and to deliver Cx43

**DOI:** 10.1101/2021.05.25.445660

**Authors:** Elena Ivanova, Carlo Corona, Cyril G. Eleftheriou, Randy F. Stout, Jakob Körbelin, Botir T. Sagdullaev

## Abstract

Endothelial cells (ECs) are key players in the development and maintenance of the vascular tree, the establishment of the blood brain barrier and control of blood flow. Disruption in ECs is an early and active component of vascular pathogenesis. However, our ability to selectively target ECs in the CNS for identification and manipulation is limited. Here, in the mouse retina, a tractable model of the CNS, we utilized a recently developed AAV-BR1 system to identify distinct classes of ECs along the vascular tree using a GFP reporter. We then developed an inducible EC-specific ectopic Connexin 43 (Cx43) expression system using AAV-BR1-CAG-DIO-Cx43-P2A-DsRed2 in combination with a mouse line carrying inducible CreERT2 in ECs. We targeted Cx43 because its loss has been implicated in microvascular impariment in numerous diseases such as diabetic retinopathy and vascular edema. GFP-labeled ECs were numerous, evenly distributed along the vascular tree and their morphology was polarized with respect to the direction of blood flow. After tamoxifen induction, ectopic Cx43 was specifically expressed in ECs. Similarly to endogenous Cx43, ectopic Cx43 was localized at the membrane contacts of ECs and it did not affect tight junction proteins. The ability to enhance gap junctions in ECs provides a precise and potentially powerful tool to treat microcirculation deficits, an early pathology in numerous diseases.

## 1 INTRODUCTION

ECs line the inside of vascular vessels and are key regulators of vascular growth (Lamalice, Le Boeuf, & Huot, 2007), homeostasis (Rubanyi, 1993), BBB (Abbott, Patabendige, Dolman, Yusof, & Begley, 2010), blood flow (Ashby & Mack, 2021; Longden et al., 2017) in health and pathological conditions. They also regulate blood fluidity (van Hinsbergh, 2012), vascular tone (Sandoo, van Zanten, Metsios, Carroll, & Kitas, 2010), and blood cell trafficking (Vestweber, 2015). To understand specific roles of ECs and target them for treatment, efficient and selective manipulation of ECs is required. Within the vascular branch, ECs tightly interact with other members of the neurovascular unit, such as pericytes and glia. These interactions are thought to be critical to vascular formation, integrity and homeostasis (Torok et al., 2021). As such, ECs can be also used as therapeutic targets to affect the function of surrounding glial cells and neurons (Y. H. Chen, Chang, & Davidson, 2009). Therefore, characterization and targeting EC sub-categories has significant yet largely untapped diagnostic and therapeutic potential.

To develop and validate a virus-mediated approach for targeting ECs we focused on the mouse retina for several reasons. The retina is a well-characterized and approachable part of the CNS. Blood vessels in the mouse retina develop after birth (Stahl et al., 2010) and are regulated by surrounding neurons (Delaey & Van De Voorde, 2000). In the retina, the vascular tree consists of distinct domains, including arterioles, vascular relays, capillaries and venules. The retina’s high metabolic demands (Wong-Riley, 2010) and the requirements for maintaining transparency of the tissue require tight control of blood flow and blood-retina barrier (BRB). Multiple vision-threatening retinal diseases have been associated with ECs. In diabetic retinopathy (DR), retinopathy of prematurity, and the wet form of age-related macular degeneration impairment of the BRB leads to leaky blood vessels and vision loss. While weakening of the BRB is typical for later stages of the disease, inadequate blood supply is present in the early stages (Patel, Rassam, Newsom, Wiek, & Kohner, 1992). Disrupted communication between vascular ECs is one of the first changes observed in diabetic retinopathy (Oku, Kodama, Sakagami, & Puro, 2001). We and others have shown that Cx43-containing gap junctions (GJs) mediate communication in healthy ECs and these GJs are selectively downregulated in ECs at early stages of DR (Bobbie et al., 2010; Ivanova, Kovacs-Oller, & Sagdullaev, 2017). Similarly to pharmacological blockade of GJs (Ivanova, Kovacs-Oller, & Sagdullaev, 2019), downregulation of Cx43 in DR leads to an impaired communication between ECs assessed by neurobiotin tracing, disrupted propagation of vasomotor response along the vascular branch, and inadequate blood flow (Kovacs-Oller, Ivanova, Bianchimano, & Sagdullaev, 2020).

Thus, Cx43 GJs in retinal ECs is a validated target for the treatment of early stages of DR. In addition, Cx43 GJs are closely co-localized and interact with tight junction proteins, including claudin 5, occludin, and ZO-1 (Ivanova et al., 2019; McCutcheon, Stout, & Spray, 2020) and participate in maintenance of barrier properties of ECs. Early rescue of Cx43 in DR may also improve later BRB disruption. As an extension of the CNS, the retina provides a good model for neuro-vascular interactions and has high unmet translational needs for targeting ECs. rAAV have been shown to have high target specificity and have been approved by the FDA for clinical use (Kuzmin et al., 2021). Multiple serotypes have been shown to transduce neurons, glia, and RPE in the retina after subretinal, intravitreal and intravenous administration (Byrne, Lin, Lee, Schaffer, & Flannery, 2015; Hickey et al., 2017). However, targeting of retinal ECs has not yet been documented.

Here we validated a newly developed AAV-BR1 viral vector for targeting ECs in the retina (Korbelin et al., 2016). First, we evaluated AAV-BR1’s transduction efficiency after intravitreal and intravenous administration using a GFP reporter. We characterized a polarized morphology of individually GFP-labeled ECs in different domains of the retinal vascular tree. Second, we developed and validated the AAV-BR1-CAG-DIO-Cx43-P2A-DsRed2-WPRE-SV40pA construct for selective and inducible overexpression of Cx43 in ECs. This vector is a valuable validated tool for regulating blood flow in the retina and potentially other parts of the CNS.

## 2 METHODS

### 2.1 Animals

All animal procedures were performed in accordance with the Institutional Animal Care and Use Committee of Weill Cornell Medicine, and in accordance with the National Institutes of Health Guide for the care and use of laboratory animals. Wildtype mice (wt) were obtained from the Jackson Laboratories (C57BL/6J, RRID:IMSR_JAX:000664). Mice expressing tamoxifen-inducible CreERT2 under the cadherin5 promoter (Cdh5CreERT2^+/−^(Sorensen, Adams, & Gossler, 2009)) were sourced from Taconic (C57BL/6-Tg(Cdh5-cre/ERT2)1Rha, RRID:IMSR_TAC:13073). Mice were bred under the fully executed breeding agreement between Dr. Sagdullaev and Taconic from 05/14/2020. Both female and male mice in equal numbers were used between p60-p100.

### 2.2 Viral constructs

AAV-BR1-CAG-GFP viral construct was designed, produced, and validated in Dr. J.Körbelin’s laboratory (Korbelin et al., 2016). In collaboration with Drs. D. C. Spray (provided rat Cx43 EBFP2-Cx43WT; (McCutcheon et al., 2020; Stout, Snapp, & Spray, 2015), we designed an EC-specific rAAV vector carrying a Cre-dependent Cx43 construct: AAV-BR1-CAG-DIO-Cx43-P2A-DsRed2-WPRE-SV40pA. The entire construct was assembled and produced by Virovek (Hayward, CA). The sequence of the construct was confirmed, and the virus was delivered to us ready for *in vivo* use.

### 2.3 Viral injections and tamoxifen treatment

For the study of AAV-BR1-CAG-GFP tropism, P60 wt mice were injected either intravitreally or intravenously. During the intravenous injection, 100 μl of the ~1 × 10^12^ GC/ml AAV-BR1-CAG-GFP construct was injected into the tail vein. For the intravitreal injections, the mice were anesthetized by intraperitoneal injection of a mixture of 150 mg/kg ketamine and 15 mg/kg xylazine. Under a dissecting microscope, a small perforation was made in the temporal sclera region with a sharp needle. A total of 1.5 μL of viral vectors suspension in saline at a concentration of ~1 × 10^12^ GC/mL AAV-BR1-CAG-GFP was injected into the intravitreal space through the perforation with a Hamilton syringe. One month after the viral injection, tissue was collected and analyzed.

For ectopic expression of Cx43, at P60 Cdh5CreERT2^+/−^ mice were injected intravenously with 100μl of the ~10^12^ GC/ml of a mixture AAV-BR1-CAG-GFP and AAV-BR1-DIO-Cx43-DsRed viral constructs. Four weeks after viral injection, one group of Cdh5CreERT2^+/−^ mice was treated with standard intraperitoneal (ip) injections of 100μl 20 mg/ml tamoxifen in corn oil daily for five consecutive days (Jackson Laboratories protocol). As a vehicle control, another group of Cdh5CreERT2^+/−^ Cre mice injected with the same virus were treated with corn oil. Three days after the final tamoxifen treatment, tissue was collected and analyzed.

### 2.4 Immunocytochemistry

As described previously (Toychiev, Sagdullaev, Yee, Ivanova, & Sagdullaev, 2013), following euthanasia, the eyes were removed and placed in bicarbonate-buffered Ames’ medium (Sigma Aldrich, St Louis, MO) equilibrated with 95% O2 and 5% CO2. The cornea was removed by an encircling cut above the ora serrata, and the iris, lens, and vitreous were extracted. The remaining eyecup, with the retina still attached to the pigment epithelium, was submersion-fixed on a shaker in freshly prepared 4% paraformaldehyde in 0.1M phosphate saline (PBS, pH = 7.3) for 20 min at room temperature. For labeling with Cx43 antibodies, the eyecups were fixed in a fixative containing: 4% carbodiimide and 0.25% paraformaldehyde in 0.1M phosphate saline (PBS, pH = 7.3) for 15 min at room temperature (Ivanova et al., 2019). After fixation, the eye cups were washed in PBS for 2 h and the retinas were detached from the eye-cups. Isolated retinas were blocked for 10 h in a PBS solution containing 5% Chemiblocker (membrane-blocking agent, Chemicon), 0.5% Triton X-100, and 0.05% sodium azide (Sigma). Primary antibodies were diluted in the same solution and applied for 72 h, followed by incubation for 48 h in the appropriate secondary antibody. In multi-labeling experiments, the tissue was incubated in a mixture of primary antibodies, followed by a mixture of secondary antibodies. All steps were completed at room temperature. After staining, the tissue was flat-mounted on a slide, ganglion cell layer up, and coverslipped using Vectashield mounting medium (H-1000, Vector Laboratories). The coverslip was sealed in place with nail polish. To avoid extensive squeezing and damage to the retina, small pieces of a broken glass cover slip (Number 1 size) were placed in the space between the slide and the coverslip. Retinal whole mounts were imaged either under a Leica SP8 confocal microscope using 20x air and 63x oil objectives or under Nikon A1R HD25 microscope using 20x water and 60x oil objectives.

All primary antibodies are listed in Table 1.

**Table 1.** Primary antibodies

| Molecular marker | Immunogen | Host | Dilution | Clone | Source | RRID |
| --- | --- | --- | --- | --- | --- | --- |
| NG2-Cy3 | Immunoaffinity purified NG2 chondroitin sulfate proteoglycan from rat | rabbit | 1:800 | poly | EMD Millipore, AB5320C3 | RRID:AB_11214368 |
| claudin5 clone 4C3C2, Alexa488 | Synthetic peptide from the mouse claudin5 | mouse | 1:10000 | mono | Invitrogen, 352588 | RRID:AB_2532189 |
| Connexin 43 (Cx43s) | PSSRASSRASSRPRPDDL EI | rabbit | 1:2000 | poly | Sigma, C6219 | RRID:AB_476857 |
| Connexin 43 (Cx43a) | (C)HAQPFDFPDDNQNSK | rabbit | 1:10000 | poly | Alomone, ACC-201 | RRID:AB_10917597 |
| Cre | synthetic peptide corresponding to residues near the | rabbit | 1:2000 | poly | Cell Signaling | RRID:AB_2798694 |
|  | amino terminus of<br>bacteriophage-P1 Cre<br>recombinase protein |  |  |  | , 15036 |  |
| Smooth muscle<br>actin, SMA,<br>1A4 | N-terminal synthetic<br>decapeptide of α-SMA | mouse | 1:1000 | mono | Sigma,<br>A5228 | RRID:AB_262054 |
| CD31/PECAM-1 | Mouse myeloma cell NS0-derived recombinant mouse<br>CD31<br>Glu18-Lys590 | goat | 1:6000 | poly | RD,<br>AF3628 | RRID:AB_2161028 |

#### SMA antibody

The antibody recognizes single isoform of α-SMA in rabbit, guinea pig, mouse, chicken, snake, sheep, goat, human, frog, rat, canine, and bovine. The antibody was raised against N-terminal synthetic decapeptide of α-SMA and tested in western blot and in immunolabeling of smooth muscle cells from bovine aortas using the monoclonal anti-ACTA2 antibody (manufacturer information). In the rat retina, the antibody labeled arterioles and the pattern was comparable with the labeling produced by two different antibodies (FITC-conjugated goat anti-mouse IgG2a (Southern Biotechnology Associates) and Texas red-conjugated sheep anti-mouse Ig (Amersham; (Hughes & Chan-Ling, 2004)).

*CD31/PECAM*-*1* Detects mouse CD31/PECAM-1 in direct ELISAs and western blots (band ~130◻kDa). It was raised against mouse myeloma cell NS0-derived recombinant mouse CD31 Glu18-Lys590 epitope. In direct ELISAs and western blots, approximately 10% cross-reactivity with recombinant human CD31 and recombinant porcine CD31 is observed. Detected mouse CD31 and rat CD31 in flow cytometry (manufacturer information). In the retina, the antibody stained vascular endothelial cells (Fang et al., 2017; Wilhelm et al., 2016).

#### Claudin5 antibody

Reactivity has been confirmed with rat, human and mouse claudin5 using rat lung, mouse kidney, mouse small intestine, mouse lung homogenates, human colon tissue, and CACO-2 human cell line. This antibody reacts specifically with the ~ 22-24 kDa endogenous claudin5 protein (manufacturer information). In the retina it labeled endothelial cells (Yanagida et al., 2017) similar to a different antibody which was validated in control and claudin5 RNAi-treated retinas (rabbit anti-claudin5, Zymed; (Campbell et al., 2009)).

#### *Connexin 43 antibody from Sigma* (Cx43s, Sigma, #C6219)

Polyclonal antibody against Cx43 was produced against a synthetic peptide corresponding to the C-terminal segment of the cytoplasmic domain (amino acids 363–382 with N-terminal added lysine) of human/rat Cx43. The antibody specificity was confirmed by Western blot and immunocytochemistry in human and rodent retina (Danesh-Meyer et al., 2012; Kerr et al., 2010). This antibody specificity was also confirmed by immunocytochemistry and western blot comparison of heart tissue from *wt* and *Cx43* KO mice (Denuc et al., 2016). This antibody is included in Validated Antibody Database (https://www.labome.com/knockout-validated-antibodies/Cx43-antibody-knockout-validation-Sigma-Aldrich-C6219.html).

#### *Connexin 43 antibody from Alomone (*Cx43a, Alomone, #ACC-201)

Polyclonal antibody against Cx43 was produced against a synthetic peptide corresponding to the C-terminal segment of the cytoplasmic domain (amino acids 331–345) of human Cx43. Rat - 14/15 amino acid residues identical; mouse - 13/15 amino acid residues identical. The antibody specificity was confirmed by Western blot in mouse brain membranes and rat heart membranes; preincubation with control peptide antigen eliminated the band (manufacturer information). In our hands, the antibody produced identical patterns in the mouse retina whole mount as a different Cx43 antibody (Sigma, #C6219, characterized above; (Ivanova et al., 2019)). These antibodies were raised against different non-overlapping epitopes. In addition, when the Alomone’s Cx43 antibody was preincubated with the control peptide antigen, the staining in the mouse retina was eliminated (Ivanova et al., 2019).

*The NG2-Cy3 polyclonal antibody* was produced by immunizing rabbits with an immunoaffinity purified NG2 chondroitin sulfate proteoglycan from rats. This antibody identifies the intact proteoglycan and core protein by WB and ELISA (manufacturer). We used this antibody as a marker of pericytes in the adult mouse retina. It labels the same vascular cells as an antibody to PDGRB (RD, AF1042) and colocalizes with cells expressing DsRed in NG2-DsRed mouse (The Jackson Laboratory, Tg(Cspg4-DsRed.T1)1Akik/J, stock #008241, RRID:IMSR_JAX:008241; (Ivanova et al., 2017)).

*The Cre recombinase monoclonal antibody* was produced by immunizing animals with a synthetic peptide corresponding to residues near the amino terminus of bacteriophage-P1 Cre recombinase protein. In Western blot analysis, the antibody recognizes Cre in transfected 293 cells but not in the control cells (manufacturer information). In our hands, the labeling was produced in inducible Cre/TdTomato mouse lines only after tamoxifen treatment but not in oil-treated Cre/TdTomato littermates. No labeling was detected in TdTomato-positive Cre-negative mice in the current study (Ivanova, Corona, Eleftheriou, Bianchimano, & Sagdullaev, 2021).

### 2.4 Statistical analysis

Statistical analysis was performed in Microsoft Excel using a t-test and ANOVA. The data are presented as mean ± standard deviation, unless otherwise indicated. The number of samples (n) indicates the number of samples per group.

## 3 RESULTS

### 3.1 Evaluation of AAV-BR1 tropism in the mouse retina after intravitreal or intravenous delivery

Specific targeting of ECs in the brain using rAAV has been previously established (Korbelin et al., 2016), however retinal delivery and tropism have not yet been assessed. Here, in the adult wt mice, we tested two experimental approaches: an intravitreal injection and a tail vein injection, previously recommended approaches for this capsid (Figure 1a). To visualize transfected cells, we used a GFP reporter under a strong ubiquitous CAG promoter (a hybrid CMV early enhancer/chicken B-actin promoter), which would potentially allow GFP expression in any retinal cell including neurons, glia and vasculature. Four weeks after the injection, GFP expression was evaluated in isolated fixed retinal whole mounts in three vascular layers and surrounding neurons (Figure 1b). To identify GFP-positive cells, the retinas were double stained for CD31 and NG2, markers for ECs and pericytes, respectively (Figure 1c,d,g,h,k,l). Following intravitreal injections, GFP-labeled cells were concentrated around the injection side throughout the entire retinal thickness (Figure 1c, g, k; right corner). Much fewer GFP-positive cells were found across the rest of the retina. We used this spatial gradient to assess concentration dependence of the viral tropism. Along the entire gradient, GFP was found in a variety of neurons including horizontal cells (Figure 1d), amacrine cells (Figure 1h) and ganglion cells (Figure 1l). No glial cells were detected. Only one EC was spotted in the center of the injection lesion (Figure 11, arrow). These findings suggest that when delivered intravitreally, the AAV-BR1 targets neurons, consistently with neuronal transduction detected in the brain (Korbelin et al., 2016).

**FIGURE 1.**
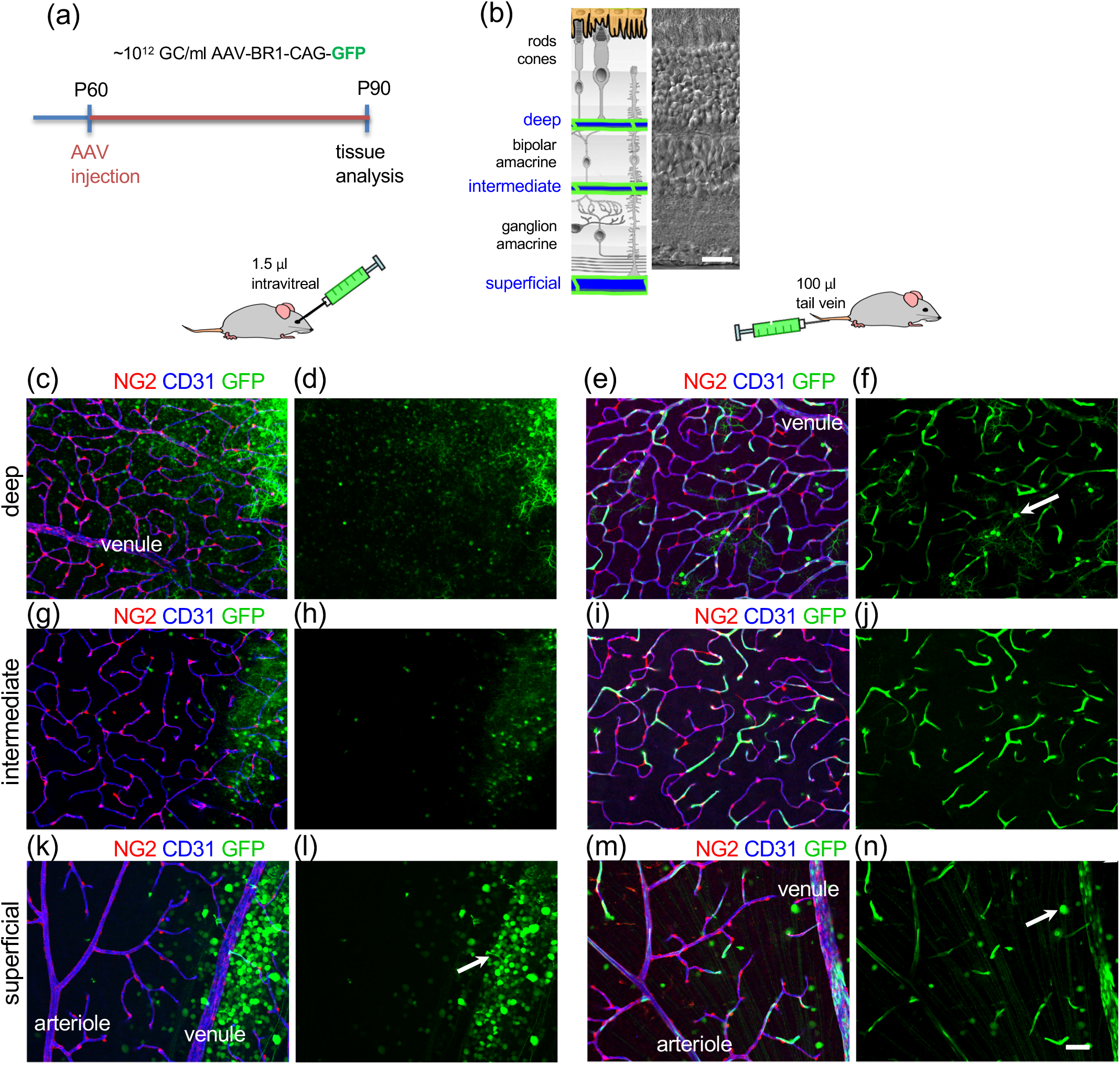
Experimental approach for AAV-BR1 injections and assessment of the virus tropism in retinal wholemounts using a GFP reporter. (a) AAV-BR1-CAG-GFP virus was delivered by either intravenous or intravitreal injections. (b) Schematic representation of retinal layers with positions of the vascular plexuses. A matched Nomarski image shows a vertical section through a mouse retina. (c-d, g-h, k-l) In a retinal wholemount after intravitreal injection (3 mice; 6 retinas), GFP was expressed in all types of neurons including photoreceptors and horizontal cells (c-d), amacrine cells (g-h), and ganglion and displaced amacrine cells (k-l). Rarely was an endothelial cell targeted (l, arrow). The majority of GFP-expressing cells were found around the injection site (right upper corner). (e-f, i-j, m-n) After intravenous injection, GFP was found predominantly in the vasculature as shown by colocalization with CD31, a marker for endothelial cells (5 mice; 10 retinas). GFP was not present in pericytes, labeled for NG2. Occasionally, single horizontal cells (f, arrow) or ganglion cells (n, arrow) were labeled. Scale bars 25 um.

On the other hand, following an intravenous injection of AAV-BR1, GFP was predominantly found in vascular cells (Figure 1e,f,i,j,m,n). All vascular layers had strongly fluorescent cells with a low number of weakly labeled neurons (horizontal cell in f, arrow and ganglion cell in n, arrow). Sporadic neuronal transduction demonstrates that AAV-BR1 can cross the BBB which was also noticed in the original study (Korbelin et al., 2016). Higher magnification images revealed a strict colocalization of GFP in the vasculature with a marker for ECs, CD31 but not with a marker for pericytes, NG2 (Figure 2a-d). Interestingly, the levels of GFP expression were highly variable even among the neighboring ECs (pairs of green and magenta arrows) and probably reflect the number of viral particles that entered the cell. GFP signals were detected in all domains of the vascular trees where it was always colocalized with ECs but not pericytes or smooth muscle cells (Figure 2e-n). In the fluorescent profiles, GFP was present in the entire cytoplasm of ECs (green line) while CD31 outlined their membrane (blue line). The ratio of GFP-labeled ECs was also similarly high in all vascular domains (Figure 2o; arteriole, 75 ± 6%; venule, 76 ± 4%; capillary, 75 ± 4%; ANOVA p=0.89; 5 retinas; 10 samples). Thus, for efficient and specific targeting of retinal ECs, AAV-BR1 must be delivered via direct injection into the bloodstream.

**FIGURE 2.**
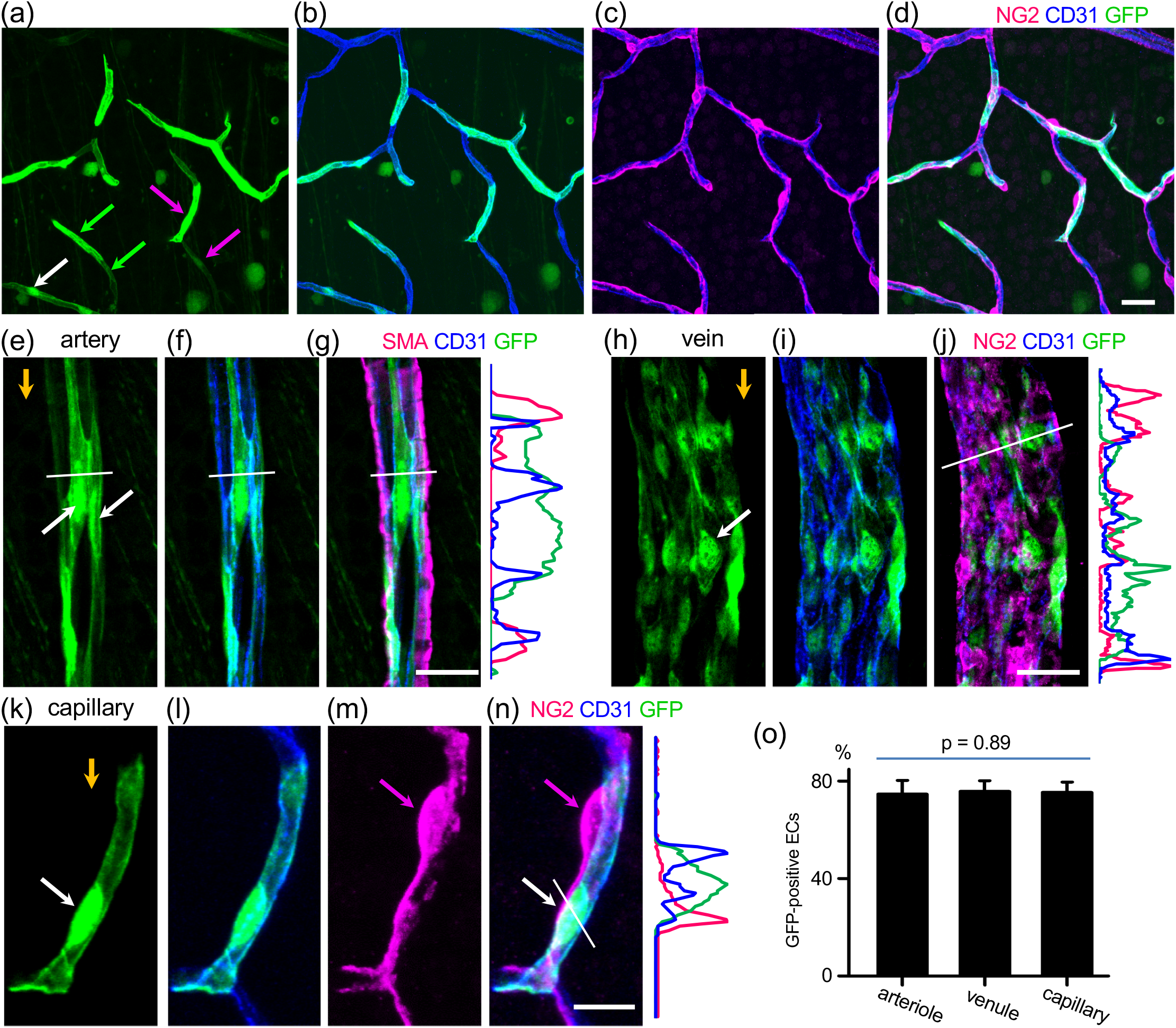
In distinct domains of the vascular tree, AAVBR1-GFP was expressed exclusively in endothelial cells but not in contractile vascular cells. (a-d) Strong variations in the levels of GFP expression was observed in ECs including neighbors (pair of green and pink arrows). No pericytes (NG2, magenta) were labeled. Within an EC, the strongest GFP fluorescence was in the nucleus (white arrow). (e-g) Multiple ECs were labeled in arterioles. Nuclei had the highest GFP fluorescence (arrows). No SMA-labeled cells presented with GFP fluorescence (SMA, magenta). Orange arrow shows the direction of blood flow. (h-j) GFP-positive ECs with brightly labeled nuclei (white arrow) in the venule. Orange arrow shows blood flow. (k-n) In the capillary, EC nuclei (white arrow) were often located close to a pericyte nuclei (magenta arrow). Scale bar 25 um in (a-j) and 10 um in (k-n). (p) The ratio of GFP-labeled ECs was similar among blood vessel types (ANOVA; 5 mice; 5 retinas; 10 samples).

### 3.2 Identification of morphological diversity and polarization of ECs

Labeling of individual ECs with AAV-BR1-GFP revealed morphologically distinct classes of ECs along the vascular tree (Figure 3a). The longest ECs were found inside the arteriole, vascular relay, and venular regions (Cells #1, 3, 8 and Figure 4b). They spanned over several contractile cells, with numerous potential sites of contact (Figure 3b, green). The length of ECs was shorter in capillaries. However, the sparse distribution of pericytes allowed for EC contact with a single pericyte (Figure 2a, cell #6). The most compact ECs were at the arteriolar branch (Figure 3a, cells # 2a). Based on the length of the ECs and distribution of contractile cells within the same regions, we estimated the number of contractile cells that can be contacted by a single EC (Figure 3b). The higher number of potential contacts (green) might indicate faster signal propagation, synchronization and higher recruitment of the contractile cells, features important for functional hyperemia. In contrast, compact ECs at the arteriolar branch may form a structural basis for their putative role as terminal cells to stop the vasomotor signal propagation and thus restricting functional hyperemia to the activated branch (Kovacs-Oller et al., 2020).

**FIGURE 3.**
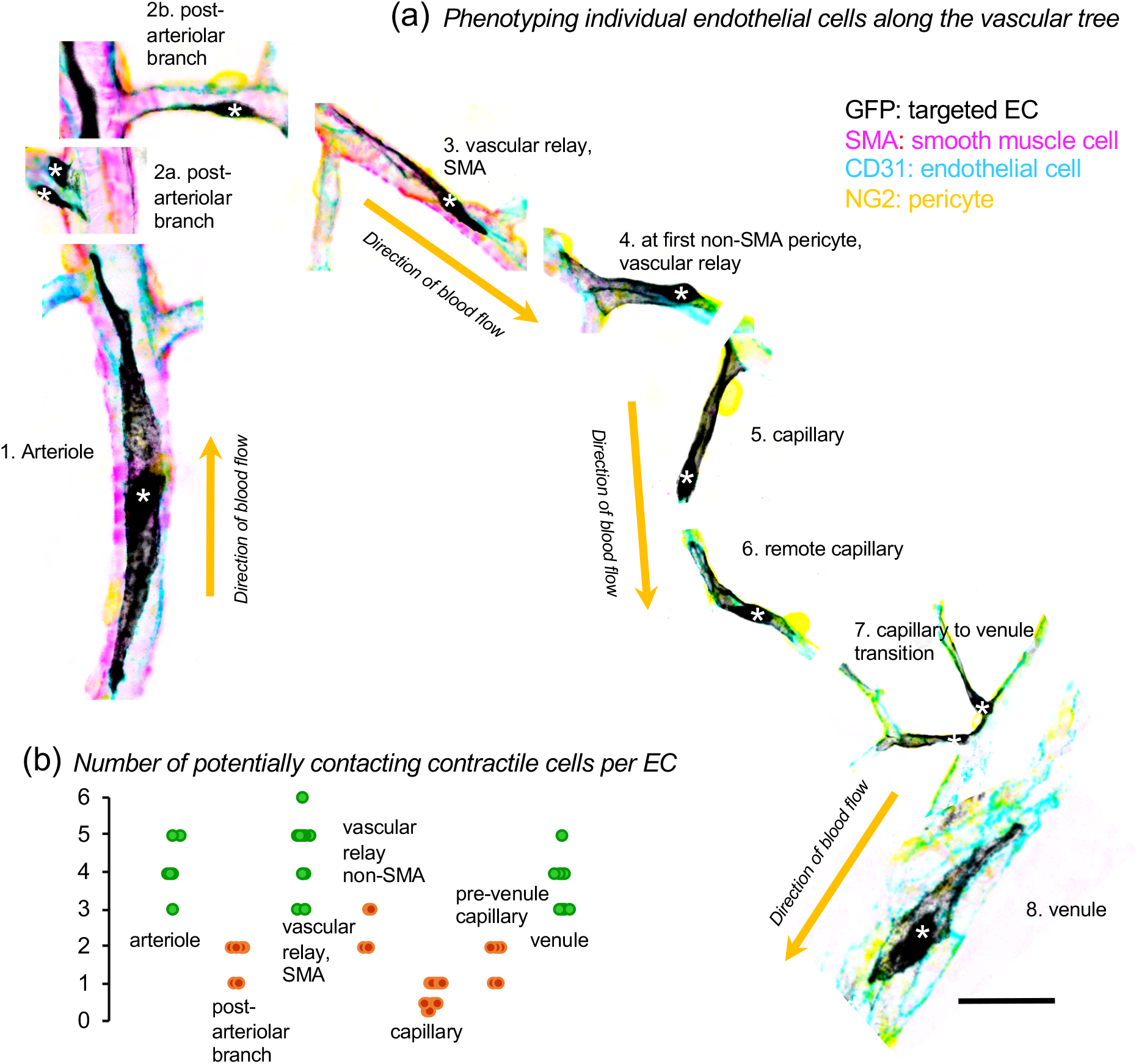
Sparse labeling reveals EC morphological diversity along the retinal vascular tree. (a) Endothelial cells were elongated along the direction of blood flow. Nuclei were shifted towards the downflow edge of cells (asterisk). At the vascular branches, numerous types of ECs were found (2a and b, 7). (b) An EC in arteriole and vascular relay spans several contractile cells; in capillaries it is limited to half of a contractile cell (usually, one pericyte spans two ECs). Scale bar 25 um.

**FIGURE 4.**
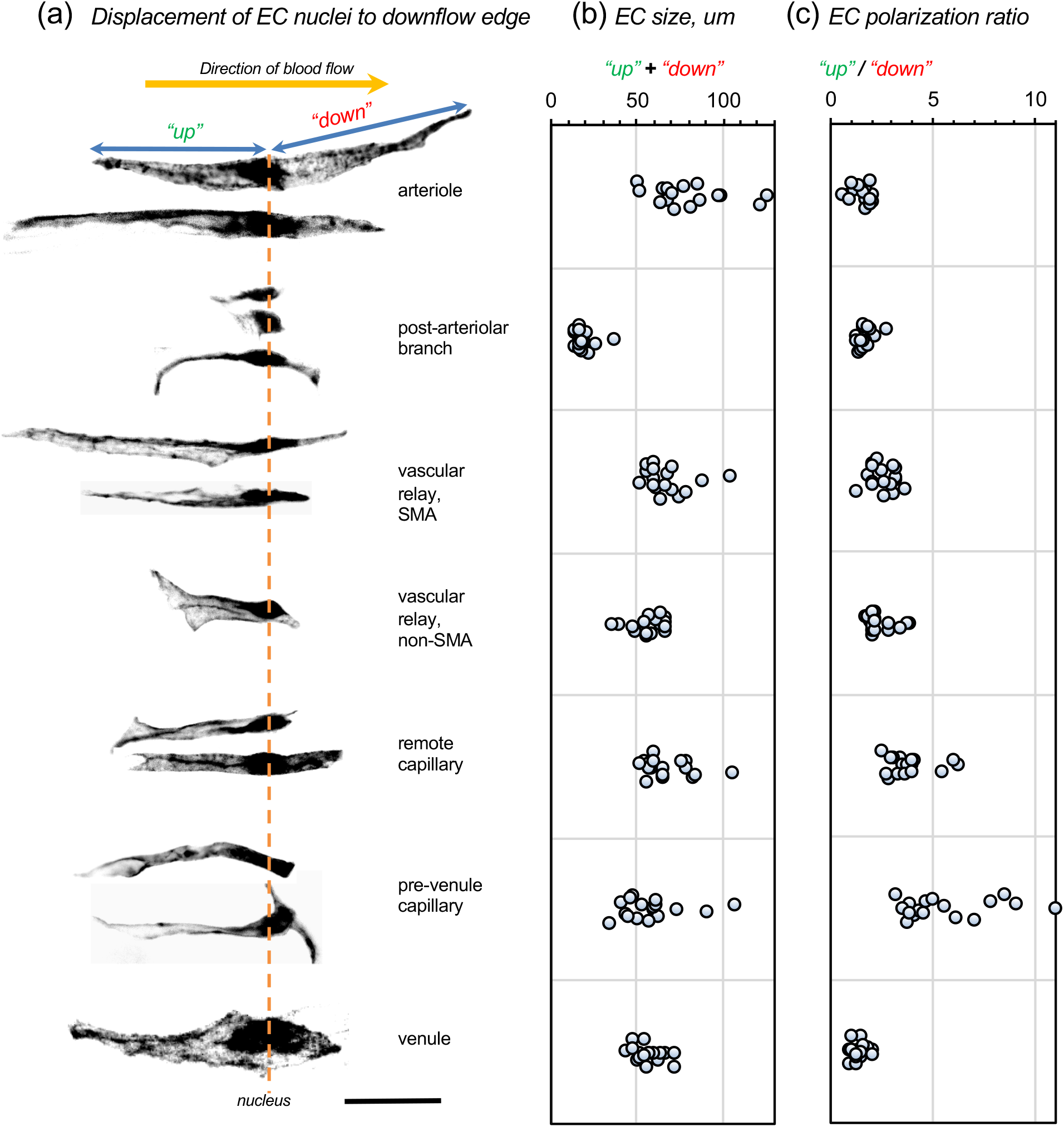
Nuclei of endothelial cells were shifted to the downflow edge. (a) ECs vertically distributed along an axis centered around their nuclei. (b) The length of ECs in distinct vascular domains. (c) EC nuclei were more displaced in narrow blood vessels (including remote and pre-venular capillaries) and were less polarized in wider arterioles and venules. 5 mice; 5 retinas; ~17 samples for each cell type. Scale bar 25 um.

Furthermore, a detailed morphometric analysis of identified ECs revealed their structural polarization (Figure 4a). In particular, ECs from the vascular relays and downstream capillaries had a pear-like shape with a long neck upstream and a wide blunt body on the downstream side. The nucleus of these ECs was shifted towards the downstream edge and the degree of the shift, designated the polarization ratio, was specific for the vascular domain. The highest ratio, indicative of the strongest shift, was found among the remote and pre-venular capillaries. The nucleus displacement was less evident in ECs of arterioles and venules (Figure 4c). In general, ECs from wide blood vessels were less polarized that the ones from narrow vessels. The prominent polarization of the ECs in capillaries might be due to elevated shear stress from blood cells crawling through narrow spaces and pushing the nuclei of ECs along their path.

### 3.3 Design and validation of a AAV-BR1-CAG-DIO-Cx43-P2A-DsRed2 to express Cx43 in retinal ECs

Using the AAV-BR1 vector and GFP reporter we established highly efficient and selective targeting of ECs in the retina. We also demonstrated a polarized morphology of ECs along the direction of blood flow. Next, we wanted to test whether the AAV-BR1 system could be utilized to selectively induce gene expression in ECs.

One of the important mechanisms controlling blood flow is the propagation of the vasomotor response, a synchronized change of the vascular diameter mediated by coordinated activity of several contractile cells in response to the stimulus. The specialized morphology and number of contacts with contractile cells can define the vasomotor response. We also previously showed that Cx43-mediated gap junctions (GJs) are essential for synchronization and propagation of the vasomotor response and, thus, control of blood flow (Ivanova et al., 2017, 2019; Kovacs-Oller et al., 2020). These GJs between ECs disappear in DR causing inadequate blood flow in the early stages (Kovacs-Oller et al., 2020) and possibly BRB impairments in later stages. To restore Cx43-containing GJs between retinal ECs, we developed an approach combining custom-made AAV-BR1-DIO-Cx43-P2A-DsRed2-WPRE-SV40 pA virus and Cdh5CreERT2^+/−^ mice (Sorensen et al., 2009). The rationale for our approach was to select specific elements, their advantages and weaknesses are summarized in Table 2.

**TABLE 2:**
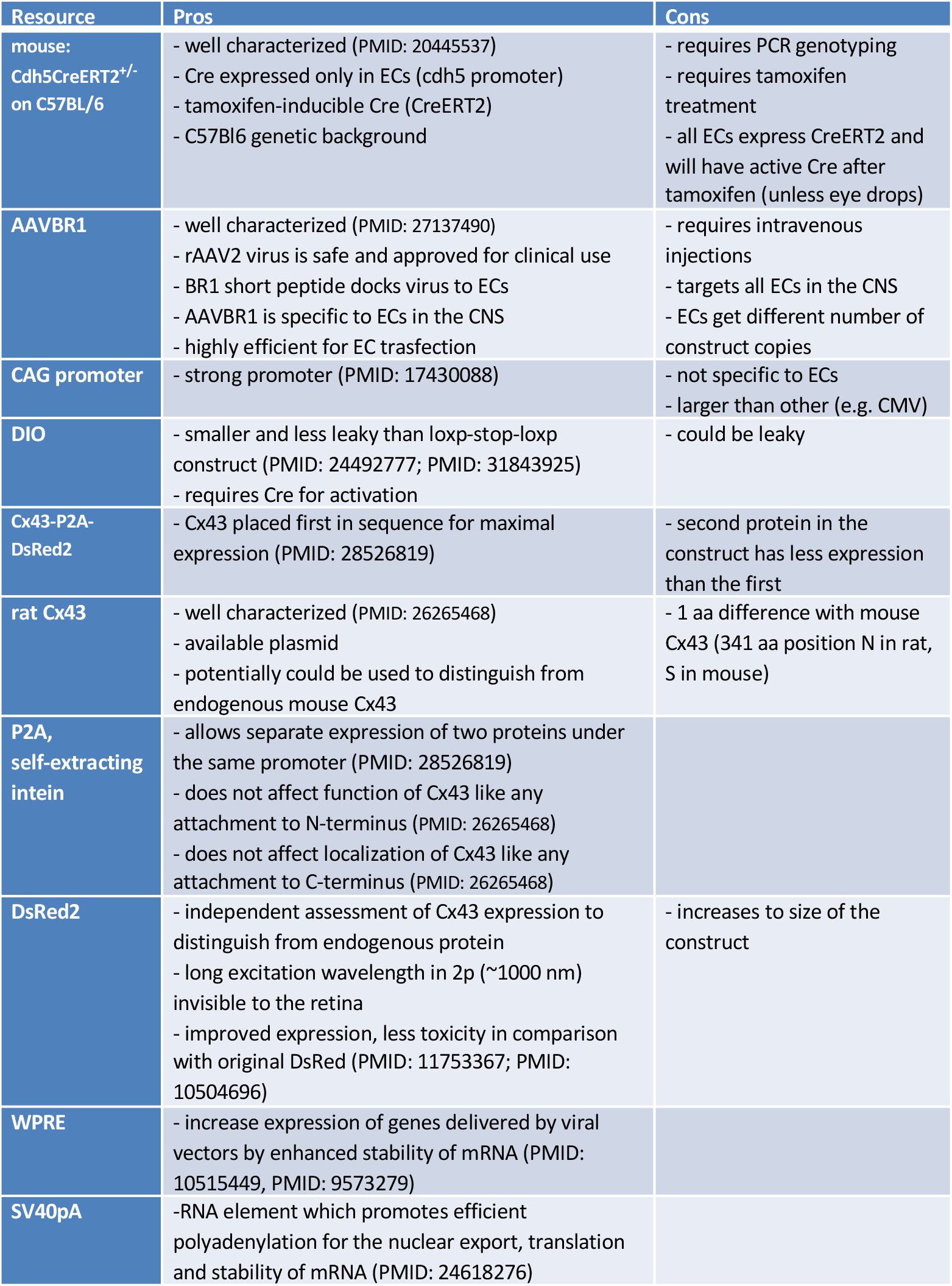
Experimental approach for controllable expression of Cx43 in endothelial cells of the mouse retina using Cdh5CreERT2+/− mice and AAV-BR1-CAG-DIO-rCx43-P2A-DsRed2-WPRESV40pA virus

| Resource | Pros | Cons |
| --- | --- | --- |
| mouse:<br>Cdh5CreERT2<sup>+/+</sup><br>on C57BL/6 | - well characterized (PMID: 20445537)<br>- Cre expressed only in ECs (cdh5 promoter)<br>- tamoxifen-inducible Cre (CreERT2)<br>- C57BL6 genetic background | - requires PCR genotyping<br>- requires tamoxifen treatment<br>- all ECs express CreERT2 and will have active Cre after tamoxifen (unless eye drops) |
| AAVBR1 | - well characterized (PMID: 27137490)<br>- rAAV2 virus is safe and approved for clinical use<br>- BR1 short peptide docks virus to ECs<br>- AAVBR1 is specific to ECs in the CNS<br>- highly efficient for EC transfection | - requires intravenous injections<br>- targets all ECs in the CNS<br>- ECs get different number of construct copies |
| CAG promoter | - strong promoter (PMID: 17430088) | - not specific to ECs<br>- larger than other (e.g. CMV) |
| DIO | - smaller and less leaky than loxp-stop-loxp construct (PMID: 24492777; PMID: 31843925)<br>- requires Cre for activation | - could be leaky |
| Cx43-P2A-DsRed2 | - Cx43 placed first in sequence for maximal expression (PMID: 28526819) | - second protein in the construct has less expression than the first |
| rat Cx43 | - well characterized (PMID: 26265468)<br>- available plasmid<br>- potentially could be used to distinguish from endogenous mouse Cx43 | - 1 aa difference with mouse Cx43 (341 aa position N in rat, S in mouse) |
| P2A, self-extracting intein | - allows separate expression of two proteins under the same promoter (PMID: 28526819)<br>- does not affect function of Cx43 like any attachment to N-terminus (PMID: 26265468)<br>- does not affect localization of Cx43 like any attachment to C-terminus (PMID: 26265468) |  |
| DsRed2 | - independent assessment of Cx43 expression to distinguish from endogenous protein<br>- long excitation wavelength in 2p (~1000 nm) invisible to the retina<br>- improved expression, less toxicity in comparison with original DsRed (PMID: 11753367; PMID: 10504696) | - increases to size of the construct |
| WPRE | - increase expression of genes delivered by viral vectors by enhanced stability of mRNA (PMID: 10515449, PMID: 9573279) |  |
| SV40pA | - RNA element which promotes efficient polyadenylation for the nuclear export, translation and stability of mRNA (PMID: 24618276) |  |

Next, we designed an EC-specific rAAV vector carrying a Cre-dependent Cx43 construct: AAV-BR1-DIO-Cx43-P2A-DsRed2-WPRE-SV40pA. We used a rat Cx43 EBFP2-Cx43WT plasmid (gift from Dr. Spray). The rat Cx43 differs from that of the mouse by one amino acid (341 aa position N in rat, S in mouse), and was chosen due to a well-characterized construct, proven to work and target Cx43 at the membrane (McCutcheon et al., 2020; Stout et al., 2015). We used a P2A self-extracting intein sequence between the Cx43 sequence and DsRed fluorescent tag. The strength of this approach is that it avoids the attachment of fluorescent tags directly to the N or C termini of Cx43, as they were shown to affect the function of the channel (N-terminus) or its incorporation and stability in the membrane (C-terminus, (Stout et al., 2015)). This results in independent synthesis of both proteins in the cell, with a stronger production of Cx43 due to its first position in the sequence (Z. Liu et al., 2017). To make the construct Cre-dependent, we created a DIO vector with the gene of interest cloned in the inverted orientation (eg. 3’-> 5’). To test for successful Cx43 expression in retinal ECs, we injected 100 μl of the ~10^12 GC/ml construct into the tail veins of Cdh5CreERT2+/− mice ((Sorensen et al., 2009), Figure 5a). In these mice, tamoxifen-inducible Cre-recombinase is expressed exclusively in ECs under the vascular endothelial cadherin (Cdh5) promoter. As a control, we co-injected the AAV-BR1-GFP virus which does not require Cre for GFP expression. Four weeks after the viral injection, the mice were treated with tamoxifen to induce Cre translocation into the nuclei (Figure 5b). No Cre was detected in the nuclei of ECs in the mouse treated with oil (Figure 5c, control).

**FIGURE 5.**
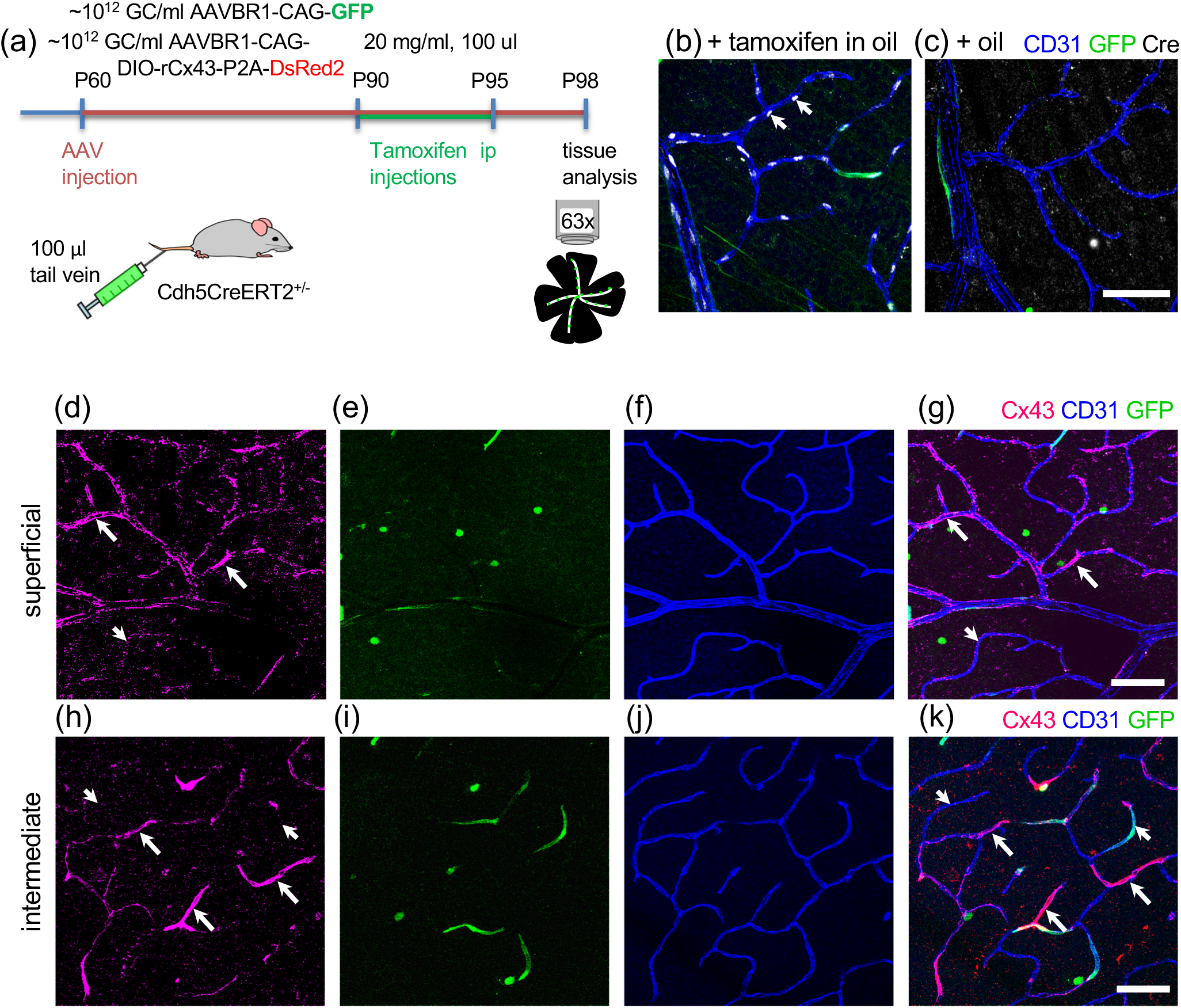
Strong and specific Cre-dependent ectopic expression of Cx43 in ECs in Cdh5CreERT2+/− mice after intravenous injection of AAV-BR1-CAG-DIO-rCx43-P2A-DsRed2. (a) Timeline and experimental procedure of AAV-mediated Cx43 expression. (b) Evaluation of Cre translocation in retinal wholemounts showed Cre in the EC nuclei of tamoxifen-treated mice (4 mice; 4 retinas). (c) Cre was not detected in oil-treated mice (3 mice; 6 retinas). (d-k) In the retina of mice injected with AAV-BR1-CAG-DIO-rCx43-P2A-DsRed2 and AAV-BR1-CAG-GFP and treated by tamoxifen, prominent Cx43 expression was found in ECs (arrows). The nearby vasculature had low endogenous Cx43 expression (arrowheads). Scale bar 50 um.

In the harvested retinal whole mounts, strong expression of ectopic Cx43 was evident in the ECs across all vascular layers (Figure 5d-k). Due to the endogenous expression of Cx43, we could not distinguish ECs with weak ectopic expression of Cx43. Nevertheless, in AAV-BR1 expressing cells, levels of Cx43 were significantly higher than in the negative neighboring ECs. Similarly to findings with AAV-BR1-GFP, ECs in all vascular domains were equally targeted.

Newly expressed Cx43 was correctly inserted into the EC membrane as confirmed by two Cx43 antibodies recognizing different epitopes (Figure 6). The high magnification images showed high concentration of Cx43 at the membrane contact of ECs, defined by CD31 labeling (arrows), similar to the localization of endogenous Cx43 (Ivanova et al., 2019). The precise localization of ectopic Cx43 to the membrane was further confirmed by fluorescent intensity profiles. As an additional control, the DsRed2 red fluorescent reporter was added to the viral construct. Unfortunately, unamplified fluorescence of DsRed2 was too low to unambiguously identify ECs transfected by the virus (Figure 6b,g,l).

**FIGURE 6.**
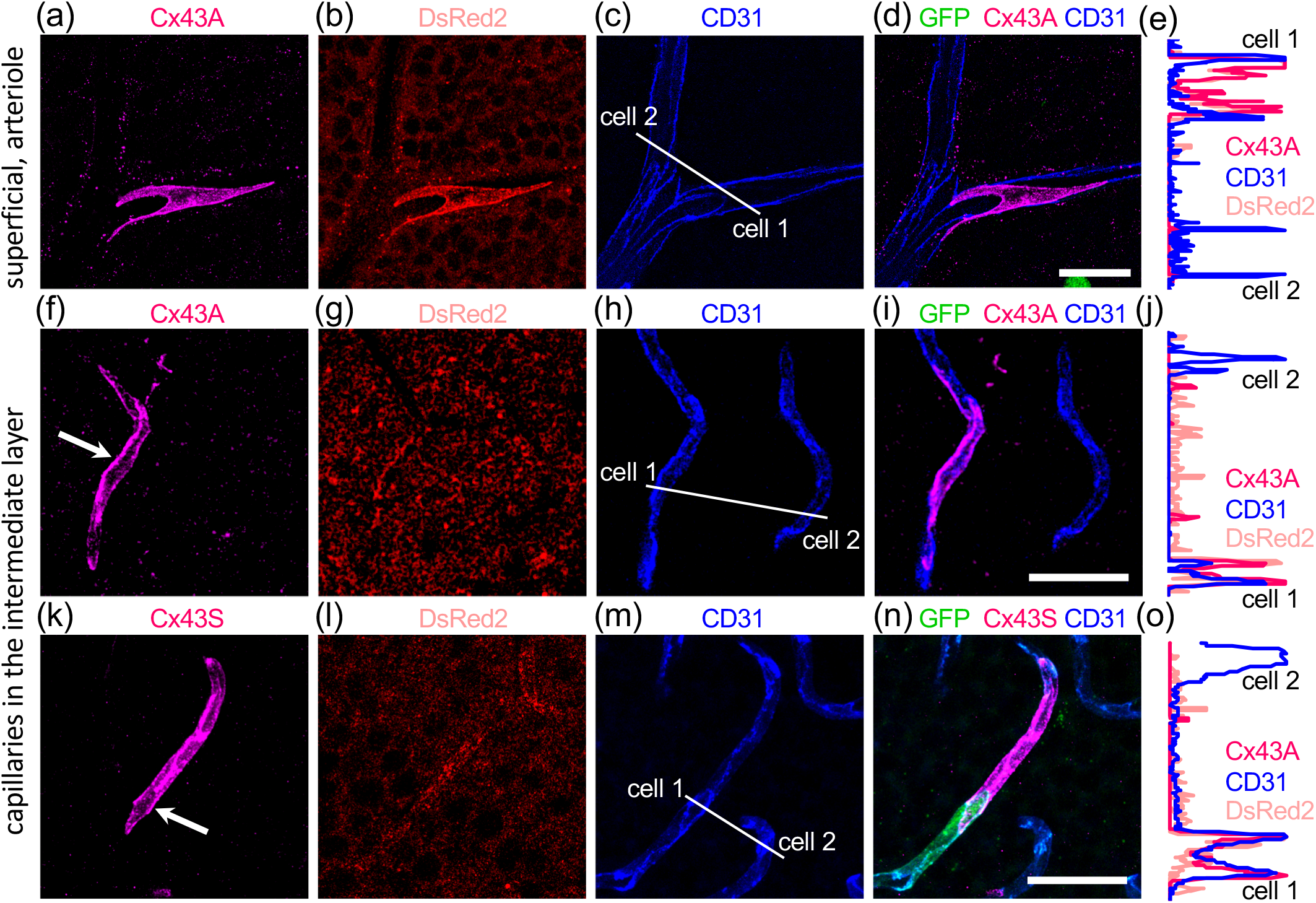
Ectopically expressed Cx43 is localized at the membrane contacts of ECs and recognized by two different antibodies. (a-j) Antibody to amino acids 331–345 from Alomone recognized ectopic Cx43. (b,g) Unamplified fluorescence of DsRed2 was very weak. (c,h) Antibody to the adherens protein CD31 was used as a marker of ECs. (d,i) Merged image. (e,j) Normalized fluorescent profile along the line in (c and h) demonstrated high membrane expression of Cx43 in cell 1. (k-o) Antibody to amino acids 363–382 from Sigma also recognized ectopic Cx43. (l) Weak expression of DsRed2. (m) Antibody to adherens protein CD31 colocalized with Cx43 labeling. (n) Merged image of (k-m). (o) Normalized fluorescent profile along the line in (m) demonstrates high membrane expression of Cx43 in cell 1. Scale bar 25 um.

In contrast to AAV-BR1-GFP, no ectopic expression of Cx43 was found in neurons. The exceptional specificity of Cx43 expression was mediated by the simultaneous requirement for Cre (specific for ECs due to cdh5 promoter), AAV-BR1 virus (specific for ECs via the intravenous route), and the requirement for Cre in DIO viral construct. No ectopic expression of Cx43 was detected in AAV-BR1-DIO-Cx43-P2A-DsRed2-WPRE-SV40pA injected Cdh5CreERT2^+/−^ mice treated by oil.

### 3.4 Selective expression of ectopic Cx43 at the membrane contacts of ECs does not interfere with tight junctions

High expression of Cx43 at the EC membrane after viral transfection may potentially affect distribution and levels of other proteins normally present at these contacts. Specifically, we were interested in whether tight junction components were affected by ectopic Cx43 expression. In double labeled ECs for Cx43 and claudin5, high levels of Cx43 did not affect distribution of claudin 5, an essential protein of tight junctions ((Nitta et al., 2003); Figure 7; compare arrow and arrowheads marked cells). This suggests that artificially high levels of Cx43 in ECs do not affect the BRB.

**Figure 7.**
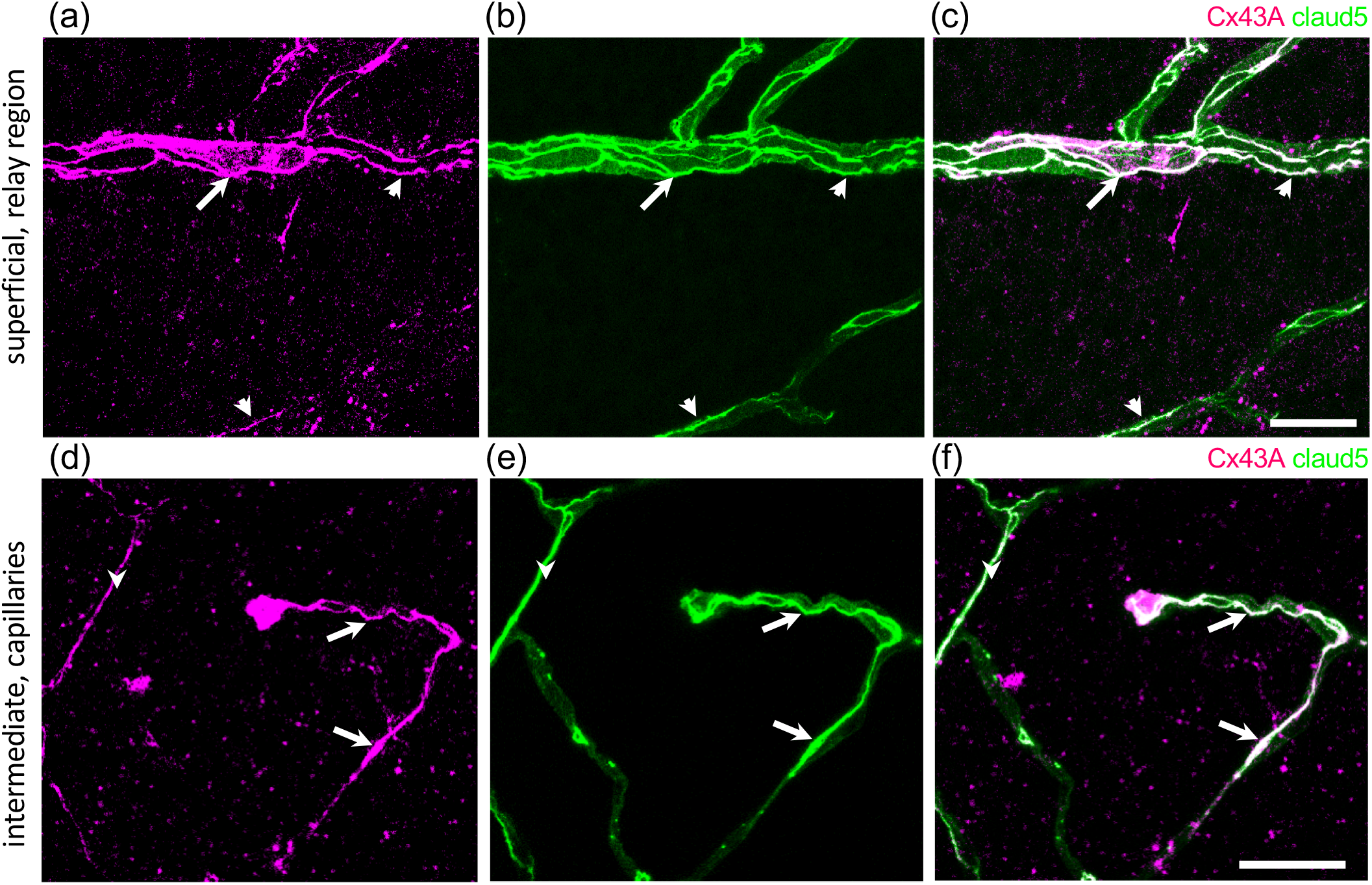
Overexpressed Cx43 does not affect localization and expression levels of claudin5. (a-f) The same location and expression levels of claudin5 were found in ECs with ectopic Cx43 (arrows) and without ectopic Cx43 (arrow heads). 4 mice; 4 retinas. Scale bars 25 um.

In conclusion, we designed and validated an efficient approach to selectively, timely, and safely express Cx43 at the membrane of ECs for control of vascular microcirculation.

## 4 DISCUSSION

The retina is one of the most metabolically demanding organs in the body and has been extensively used as a model system to study CNS angiogenesis and blood flow. Here, we developed an approach to selectively label and manipulate retinal endothelial cells (ECs), the building and functional foundation of the vasculature. We used a combinatorial approach with an EC-specific AAV-BR1 virus and a mouse carrying an inducible Cre under the cdh5 promoter. We found that this virus is equally effective for ECs in the entire retinal vascular tree. Across distinct vascular domains, individually GFP-labeled ECs exhibited different shapes, sizes and polarizations along the blood flow and number of potential contacts with contractile cells. Our custom made AAV-BR1-CAG-DIO-Cx43-P2A-DsRed2 virus allowed targeted Cx43 expression in ECs. Similarly to endogenous Cx43, ectopic Cx43 was localized to the membrane contacts of the ECs while maintaining other adherens and tight junction proteins. Below, we discuss the main findings, rational and potential for targeting ECs.

### 4.1 ECs diversity and blood flow

ECs are the building blocks of all blood vessels and play an essential role in many physiological processes in health and disease. These include neovascularization and angiogenesis, control of blood flow, vascular tone, BBB and hemostatic balance (Aird, 2007). Despite a common terminology, ECs represent diverse and functionally unique classes of cells (Aird, 2012; Kalucka et al., 2020). They differ by their tissue-specific roles, location in the vascular domain, morphology and expression of cell-specific markers. Each EC class is tailored to meet the demands of the underlying tissue (Jambusaria et al., 2020).

While ECs are actively involved in the regulation of blood flow, they are also shaped by the blood flow through polarization and protein expression. Shear stress, a frictional force acting in the direction of blood flow on the surface of ECs, and pressure◻stretch which acts perpendicularly to the vascular wall affect ECs. Increased shear stress stimulates the release of EDRF and PGI2 probably via activation of a K+ channel (inward rectifier) in the EC membrane (Rubanyi, Freay, Kauser, Johns, & Harder, 1990). In cell culture, ECs which have a polygonal shape at rest become gradually oriented and elongated in the direction of flow upon increasing shear stress (Malek & Izumo, 1996). This reorientation streamlines the ECs, decreasing the effective resistance, which may be of importance in terms of adaptation to shear stress stimuli. In *in vivo* studies, the nuclear pattern of ECs in canine arterial vessel segments was realigned in the direction of blood flow within 10 days after flipping of the segments (Flaherty et al., 1972). In the current study, nuclear polarization and shapes of ECs were evaluated in the entire retinal vascular tree in vivo. We found a prominent polarization of the nucleus, and a narrow shape of the ECs with the strongest effect in narrow blood vessels. We also found that the number of contractile cells contacting a single ECs was the greatest in vascular relays, venules, and arterioles, the blood vessels known to produce the strongest and fastest vasomotor response (Hartmann et al., 2021). In contrast, capillaries, with their relatively limited ability for vasomotor response had fewer contacts. The conspicuously low number of contacts at branching points, along with the compact shape of ECs at this location may provide an explanation for our previous work, documenting a termination of the vasomotor signal at vascular branches (Kovacs-Oller et al., 2020; Zhang, Wu, Xu, & Puro, 2011). This termination is essential for isolating active vascular branches of the vascular tree and localizing functional hyperemia (Kovacs-Oller et al., 2020).

### 4.2 Signaling between ECs and the role of Cx43

ECs regulate blood flow by influencing vascular contractile cells using both vasoactive messengers and direct GJ-mediated contacts. The vasoactive agents include nitric oxide, endothelium-derived hyperpolarizing factor, eicosanoids and endothelin-1 (Cohen, 1995). Endothelium-derived hyperpolarizing factor hyperpolarizes the vascular smooth muscle to induce relaxation of the vessel wall (Feletou & Vanhoutte, 2006). NO production by ECs is constitutive but may be regulated (Tousoulis, Kampoli, Tentolouris, Papageorgiou, & Stefanadis, 2012). Endothelin is a known vasoconstrictor secreted by the ECs that can reduce local cerebral blood flow (Dhaun & Webb, 2019). Blood flow regulation requires propagation of the vasomotor signal upstream the vascular branch. The coordinated activity of ECs and contractile vascular cells is largely mediated by GJs (B. R. Chen, Kozberg, Bouchard, Shaik, & Hillman, 2014; Hartmann et al., 2021; Kovacs-Oller et al., 2020; Longden et al., 2017). In some experiments, the damage to ECs along the vascular branch prevented upstream propagation of the signal (B. R. Chen et al., 2014). In particular, in a mouse model of diabetic retinopathy, Cx43 GJs of ECs along the vascular relay were diminished early in the disease (Ivanova et al., 2017). Their disappearance was shown to impair vasomotor signal propagation and to affect blood flow (Ivanova et al., 2017; Kovacs-Oller et al., 2020). ECs also synchronize contractile vascular cells (Hartmann et al., 2021). Densely packed and electrically coupled contractile cells on a feeding arteriole provide a much stronger and faster response than a single pericyte straddling a capillary vessel (B. R. Chen et al., 2014; Emerson & Segal, 2000; Hartmann et al., 2021; Iadecola, Yang, Ebner, & Chen, 1997; Kovacs-Oller et al., 2020; Longden et al., 2017).

In our previous work, we identified and characterized the GJs responsible for regulation of the vasomotor response and blood flow. These are unique string-like GJs of ECs in the vascular relay, a micro-vascular circuit interfacing capillaries to the feeding arteriole, interconnected by Cx43-containing gap junctions (GJs) along the contacts of endothelial cells (Ivanova et al., 2019; Kovacs-Oller et al., 2020). Cx43 GJs were essential for GCamp6f signal propagation, vasomotor response propagation, and tracer coupling (Ivanova et al., 2019; Kovacs-Oller et al., 2020). Distribution of Cx43 along the vascular relay stops abruptly at the vascular branch as shown by neurobitin tracing (Kovacs-Oller et al., 2020), immunocytochemistry (Ivanova et al., 2019) and physiological measurements (Ivanova et al., 2019; Kovacs-Oller et al., 2020; Zhang et al., 2011) where ECs are connected by Cx40 (Ivanova et al., 2019; Zechariah et al., 2020). Thus, GJs and specifically Cx43 of the vascular relay are involved in the regulation of blood flow. Their control is essential for basic and clinical studies and can be accomplished by our viral approach.

### 4.3 Implication of targeting ECs for control of microvasculature

Currently, AAV vectors have been proven to provide required specificity, efficiency, and safety in basic science and clinical applications (Kuzmin et al., 2021; Naso, Tomkowicz, Perry, & Strohl, 2017; Young, Searle, Onion, & Mautner, 2006). The initial development of the AAV-BR1 virus specific for ECs in the CNS (Korbelin et al., 2016) opened a vast field of opportunities for unmet basic and clinical applications. Due to its high specificity and efficiency, this vector allowed for repairing the BBB in *incontinentia pigmenti* disease, where a deficiency of the Nemo gene leads to a loss of brain ECs and a breakdown of the BBB (Dogbevia et al., 2017; Korbelin et al., 2016). Transduction of CNS endothelial cells with the AAV◻BR1 vector could potentially be tuned to improve cerebral perfusion after a stroke (Park et al., 2021), impair vascularization of brain tumors, repair or otherwise modulate the BBB (Santisteban et al., 2020), and directly target the CNS through systemic blood circulation (epilepsy, (X. X. Liu et al., 2020); Alzheimer’s, (Zou et al., 2020)).

This virus could be utilized to dissect the interactions between ECs and surrounding neurons. It was used to demonstrate hippocampal-dependent memory and reduced dendritic spine density in CA1 neurons in mice after selective knockout of semaphorin 3G in ECs (Tan et al., 2019); it was further used to dissect the role of semaphorin 3G in ischemic retinopathy (D. Y. Chen et al., 2021). ECs could also be targeted for the delivery of proteins to underlying neural cells (D. Y. Chen et al., 2021). In the current study, we used the virus to selectively enhance Cx43 expression in ECs which have previously been shown to be involved in the regulation of blood flow (Figueroa & Duling, 2009; Pohl, 2020; Puro, 2012) and are downregulated in the early stages of diseases like diabetic retinopathy (Ivanova et al., 2017; Kovacs-Oller et al., 2020). GJs may be involved in the regulation of barrier functions (Kandasamy, Escue, Manna, Adebiyi, & Parthasarathi, 2015; Li, Mruk, Lee, & Cheng, 2010; McCutcheon et al., 2020; Nagasawa et al., 2006; Tien, Muto, Barrette, Challyandra, & Roy, 2014). Thus, a tool to selectively manipulate GJs in ECs is important for basic and clinical applications.

The AAV-BR1 system will undoubtedly enable significant import in targeting ECs. The AAV-BR1 system will undoubtedly enable significant import in targeting ECs. However, further improvements will be needed. Capsid modifications and specific promoters might be needed to restrict the expression to specific domains of the vascular tree. The system would also benefit from modifications to avoid the spleen as most AAV are trapped there (Korbelin et al., 2016; Zhao et al., 2020). Also some modifications are required to escape from neutralizing antibodies (Bello et al., 2009; Lugin, Lee, & Kwon, 2020; Tse et al., 2017). In addition, vectors specific for pathological states are required to spare healthy cells from treatment (Guenther et al., 2019).

## ACKNOWLEDGMENTS

This work was supported by NIH grants R01-EY026576, R01-EY029796 and NYSDOH-C34457GG (BTS). The authors would like to acknowledge the Structural and Functional Imaging Core at Burke Neurological Institute and NIH S10 shared Instrumentation Grant OD028547-01 for supporting this work. The authors are grateful to Dr. David Spray for a gift of the rat Cx43 EBFP2-Cx43WT plasmid.

## AUTHOR CONTRIBUTIONS

EI, BTS, CC, and CGE conceived and performed the experiments and analysis. RFS, JK provided the Cx43 plasmid and AAV-BR1 and participated in the design of the AAV-BR1Cx43 construct. All authors contributed to writing the paper.

## DATA AVAILABILITY STATEMENT

The data that support the findings of this study are available from the corresponding author/s upon reasonable request.

## Notes

**Funding statement:** Supported by NIH grants R01-EY026576, R01-EY029796 and NYSDOH-C34457GG (BTS).

**Conflict of interest disclosure:** The authors declare no potential conflict of interest. Jakob Körbelin is listed as inventor of a patent on AAV-BR1 (#10696717) held by Boehringer Ingelheim International and serves as scientific advisor on the topic of AAV vector development.

### Competing Interest Statement

The authors declare no potential conflict of interest. Jakob Korbelin is listed as inventor of a patent on AAV-BR1 (#10696717) held by Boehringer Ingelheim International and serves as scientific advisor on the topic of AAV vector development.

